# Visualization of the initial fixed nitrogen transport in nodulated soybean plant using [^13^N]N_2_ tracer gas in real-time

**DOI:** 10.1101/2020.03.24.006049

**Authors:** Hung V. P. Nguyen, Hue Tong Thi

**Affiliations:** Institute of Biotechnology, Hue University, Vietnam

**Keywords:** ^13^N_2_, positron-emitting tracer imaging system, nitrogen fixation, nodule, soybean, transport

## Abstract

The observation of initial transport of fixed nitrogen in intact soybean plants in real-time was conducted by using the positron-emitting tracer imaging system (PETIS). Soybean root nodules were fed with [^13^N]N_2_ for 10 minutes, and the radioactivity of [^13^N]N tracer was recorded for 60 minutes. The serial images of nitrogen fixation activity and translocation of fixed nitrogen in the soybean plant were reconstructed to estimate the fixed-N transport to the upper shoot. As a result, the signal of nitrogen radiotracer moving upward through the intact stem was successfully observed. This is the first report that the translocation of fixed-N is visualized in real-time in soybean plant by a moving image. The signal of nitrogen radiotracer appeared at the base stem at about 20 minutes after the feeding of tracer gas and it took 40 minutes to reach the upper stem. The velocity of fixed nitrogen translocation was estimated approximately at 1.63 cm min^-1^. The autoradiography taken after PETIS experiment showed a clear picture of transport of fixed ^13^N in the whole plant that the fixed-N moved not only via xylem system but also via the phloem system to the shoot after transferring from xylem to phloem in the stem although it has been generally considered that the fixed-N in nodule is transported dominantly via xylem by transpiration stream toward mature leaves. This result also suggests that the initial transport of fixed-N was mainly into the stem and subsequently translocated to young leaves and buds via the phloem system. These new findings in the initial transport of fixed nitrogen of soybean by PETIS observation will become the basis for future study of fixed-N transport in the whole legume plants.

## Introduction

Biological nitrogen fixation (BNF) is very important not only for plant life especially in legume plants but also for nitrogen cycling of the globe. One of the most important characters of the legume plant is that it can use N_2_ gas in the atmosphere as a nutrient source for growth and development via biological fixation pathway in symbiotic with bacteroids in root nodules. Since, the understanding of biological nitrogen fixation and fixed nitrogen transport is very important for applying to legume cultivation to increase crop productivity, so that the dynamic of BNF and fixed-N transport have been concerning many researchers.

In soybean plant, after fixation, a major part of fixed-N is metabolized to ureides in nodules and then transported to the upper parts including shoots, leaves, and pods via xylem system and redistributed to pods, seeds, and roots via phloem system (Ohyama et al., 2009). There are two main routes of fixed-N transport in legume plants. One, the fixed-N is moved from root nodules via the xylem system to shoot. The other one, the fixed-N after incorporating into various N compounds in mature leaves moved from the mature leaves to growth organs or the storage organs by the phloem system (Oghoghorie and Page, 1972).

Up to now, various methods have been used in the field of BFN, of which the positron-emitting tracer imaging system (PETIS) developed in recent decades for researching in the field of plant nutrition is considered one of the most advanced method. This method can overcome the obstacle that previous methods could not perform. Especially, the PETIS system can detect γ-ray created by positron-emitting nuclide and can observe the movement of labeled elements in living plants in real-time (Kume et al., 1997). This technique can visualize the dynamic transport and allocation of metabolites at large distance scales and give information for understanding the whole plant’s physiological response to environmental changes in real-time (Kiser et al., 2008). PETIS was used successfully in the first real-time images of nitrogen fixation activity in an intact soybean plant (Ishii et al., 2009), and in the analysis of nitrate transport in soybean (Sato et al., 1999). Recently, by applying mathematical models in quantifying of radioisotope activity in time course, the rate import and export of radioactive tracers were calculated relatively accurate in broad bean (Matsuhashi et al., 2005), soybean (Ishii et al., 2009).

In this study, PETIS was used to elucidate more clearly the pathway that fixed nitrogen was transported and translocated in soybean plants in real-time.

## Materials and Methods

### Plant materials and cultures

Soybean (*Glycine max* [L.]Merr. cv. Williams) seeds were sterilized with 70% ethanol for 30 seconds and sodium hypochlorite solution 0.5% for 5 min and then thoroughly washed with deionized water. The seeds were inoculated with the suspension of *Bradyrhizobium japonicum* (strain USDA 110) and sown on a vermiculite tray. Ten days after sowing, the seedlings were transferred to plastic containers containing 20 L of nitrogen-free nutrient solution (K_2_SO_4_:109 mgL^-1^, K_2_HPO_4_: 8.5mg L^-1^, KCl: 0.935 mg L^-1^, CaCl_2._H_2_O: 183.0 mg L^-1^, MgSO_4._7H_2_O: 123 mg L^-1^, H_3_BO_4:_ 0.367 mg L^-1^, CuSO_4._5H_2_O: 0.032 mg L^-1^, MnSO_4_ 0.189 mg L^-1^, ZnSO_4._7H_2_O: 0.144 mg L^-1^, NiSO_4._6H_2_O: 0.0035 mg L^-1^, ethylenediamine-tetraacetic acid.2Na: 18.6 mg L^-1^, FeSO_4_.7H_2_O: 13.9 mg L^-1^; pH: 6.0). The new solution was changed every week and aerated by a pump system. Soybean plants were cultivated in a growth chamber under the conditions of 16 h light at 28^°^C and 8 h darkness at 18^°^C; humidity: 65%; and irradiance: 400 μE m-1 x s-1 under florescence light-tubes. Twenty-five to thirty day-old plants were used for the ^13^N experiments.

### Synthesis of [^13^N]N_2_ gas

The [^13^N]N_2_ was produced at the cyclotron facilities at TIARA (Japan Atomic Energy Agency, Takasaki, Gunma, Japan) by bombarding CO_2_ for ten minutes with 0.5µA of 18.3 MeV proton beam delivered from a cyclotron.

The rapid production method of the [^13^N]N_2_ tracer based on a previous study (Ishii et al., 2009) and some modification was described as follows: 38 mL of pure CO_2_ gas was filled into a target chamber with 5×10^5^ Pa and then it was irradiated with a proton beam delivered from a cyclotron. After irradiating, 15 mL of non-radioactive nitrogen gas was added to the target chamber as the carrier gas in order to carry the N radioactive from the target chamber to the receiver. The mixed gases after irradiating (including CO_2_, [^13^N]N_2_, [^13^N]N_2_O and N_2_) were purified by passing through a glass column containing soda lime powder (Soda-lime No.1; Wako Pure Chemical Industries, Osaka, Japan) to absorb all CO_2_, and then mixed gas went through a glass column containing pure granular copper (LUDISWISS, Switzerland) placed in a furnace at 600^°^C to deoxidize [^13^N]N_2_O to [^13^N]N_2_. The purified gas was collected in a syringe for checking contamination by gas chromatography. After purifying, 25 mL of the pure [^13^N]N_2_ gas was mixed 15 mL non-radioactive nitrogen and 10 mL O_2_ gas to make the final composition of O_2_: N_2_ = 2:8 for the experimental treatment.

### [^13^N]N_2_ tracer gas treatment and imaging with PETIS

Soybean plants at 26-day-old were fed with [^13^N]N_2_ gas in the PETIS system at TIARA (Japan Atomic Energy Agency, Takasaki, Gunma, Japan). The PETIS system for imaging experiment was set up as shown in **figure 1**. The root system of soybean plants was inserted into an acrylic box and the base stem of the plant at the top of the acrylic box was sealed by plastic clay to prevent gas leakage. The inlet and outlet of the gases and solution were connected with silicon tubes and controlled by valves (**Figure 1A**). The acrylic box was placed in the middle between the two detector heads of PETIS (Modified type of PPIS-4800; Hamamatsu Photonics, Hamamatsu, Japan) in a growth chamber (**Figure 1B**) with relative humidity of 65% at 28°C so that the main observation area was located at the center of the field of view (FOV). The light was maintained at a photon flux density of approximately 150 µmol photon m^-2^s^-1^.

**Figure 1:**
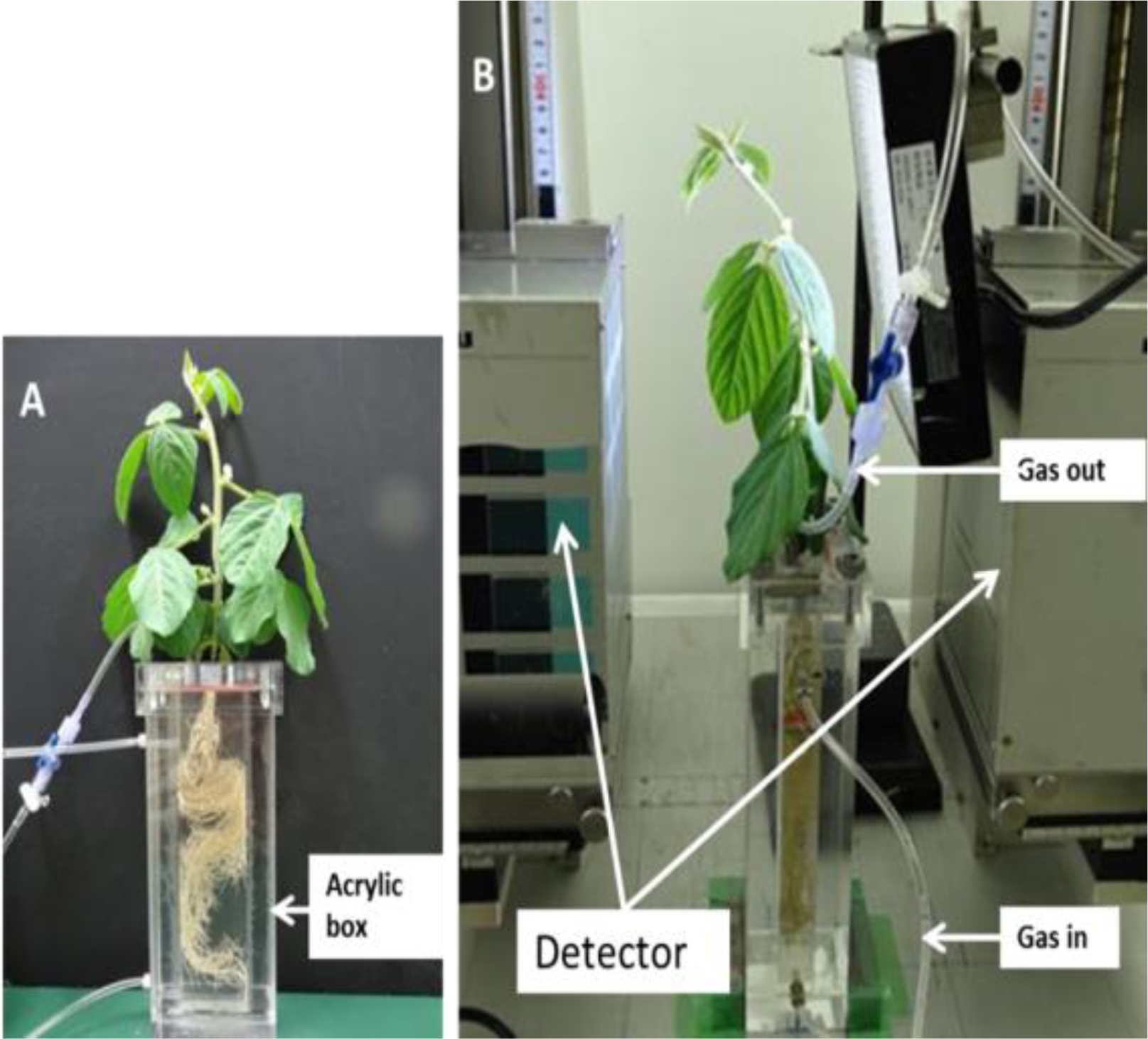
Soybean plant was set up for [^13^N]N_2_ experiment. A: The test soybean plant in an acrylic box. B: PETIS imaging System

The [^13^N]N_2_ gas treatment was implemented as following steps: First, the root system of soybean plants was adapted to a non-radioactive gas for 30 min, and then the culture solution in the acrylic box was raised to the inner top of the acrylic box to flush out the initial gas. Subsequently, 50 mL of solution was drained off and 50 mL of the fed gas containing [^13^N]N_2_ was introduced to the box at the same time. The ^13^N tracer gas was kept for 10 min in the acrylic box and flushed out by raising the solution in the acrylic box.

PETIS imaging was started when [^13^N]N_2_ tracer was filled in the acrylic box. Each frame (image) was obtained every 10 seconds for 1 hour.

### Estimation of nitrogen fixation rates and fixed N transport rate

All PETIS image data were reconstructed and analyzed by using NIH image J 1.45 software. To estimate the dynamic of [^13^N]N_2_ accumulated in nodules and the translocation of fixed [^13^N]N_2_ from nodules to the upper stem, the regions of interest (ROIs) on the integrated PETIS images were drawn and extracted at clump of nodules and the time-activity curves (TACs) were created from ROIs of a series PETIS images in 60 minutes, these curves were corrected for physical decay of [^13^N]N_2_. The data of TACs will be used for estimating the rate import and export of [^13^N]N_2_ at ROIs.

To estimate the fixed-N activity, the average radioactivity (Bq) of the first 10 frames after the flushing out of [^13^N]N_2_ tracer gas was calculated and then converted into the amount of total nitrogen (µmol N_2_). This value indicates the amount of total nitrogen fixed by the nodules during 10 minutes of ^13^N exposure and was used to estimate the rate of nitrogen fixation (µmol N_2_ h^-1^).

To analyze the export of fixed nitrogen from the nodules, a linear regression was made on the data points of the time-activity curve of each sample for 20 minutes from the end of flushing out of the tracer gas, and the slope of the line was converted to the decreasing rate of fixed N in nodule (µmol N_2_ h^-1^).

#### BAS imaging

To obtain autoradiography images, the plants were exposed to the imaging plates of a bio-imaging analyzer (BAS GUGE 2040, Fujifilm, Tokyo, Japan) for 30 minutes. After exposure, the plates were scanned with a bio-imaging analyzer system (GE Healthcare, Typhoon FLA 7000). The autoradiograph image was reconstructed by using the NIH image J1.45.

## Results

In the [^13^N]N_2_ tracer experiment using the PETIS system, the nitrogen fixation activity and the transport of fixed nitrogen are reflected by the [^13^N]N_2_ radioactivity accumulated at a detected area of the plant. **Figure 2** shows the test plant and serial images recorded by PETIS after flushing out of [^13^N]N_2_ gas. Due to the small FOV, the detectors were only concentrated around the upper nodules and lower shoot (**Figure 2A**) to observe the dynamics of nitrogen fixation in the nodules and transport of the fixed-N to upperparts. The PETIS images were taken every ten seconds, and **figure 2B** shows the restacked images in a sequence of 5 minutes (equal to 30 frames) of all frames. It was demonstrated that just after five minutes exposing to [^13^N]N_2_ gas, the fixed-N has already been at the base stem and then gradually moved up to shoot (**Figure 2B**).

**Figure 2:**
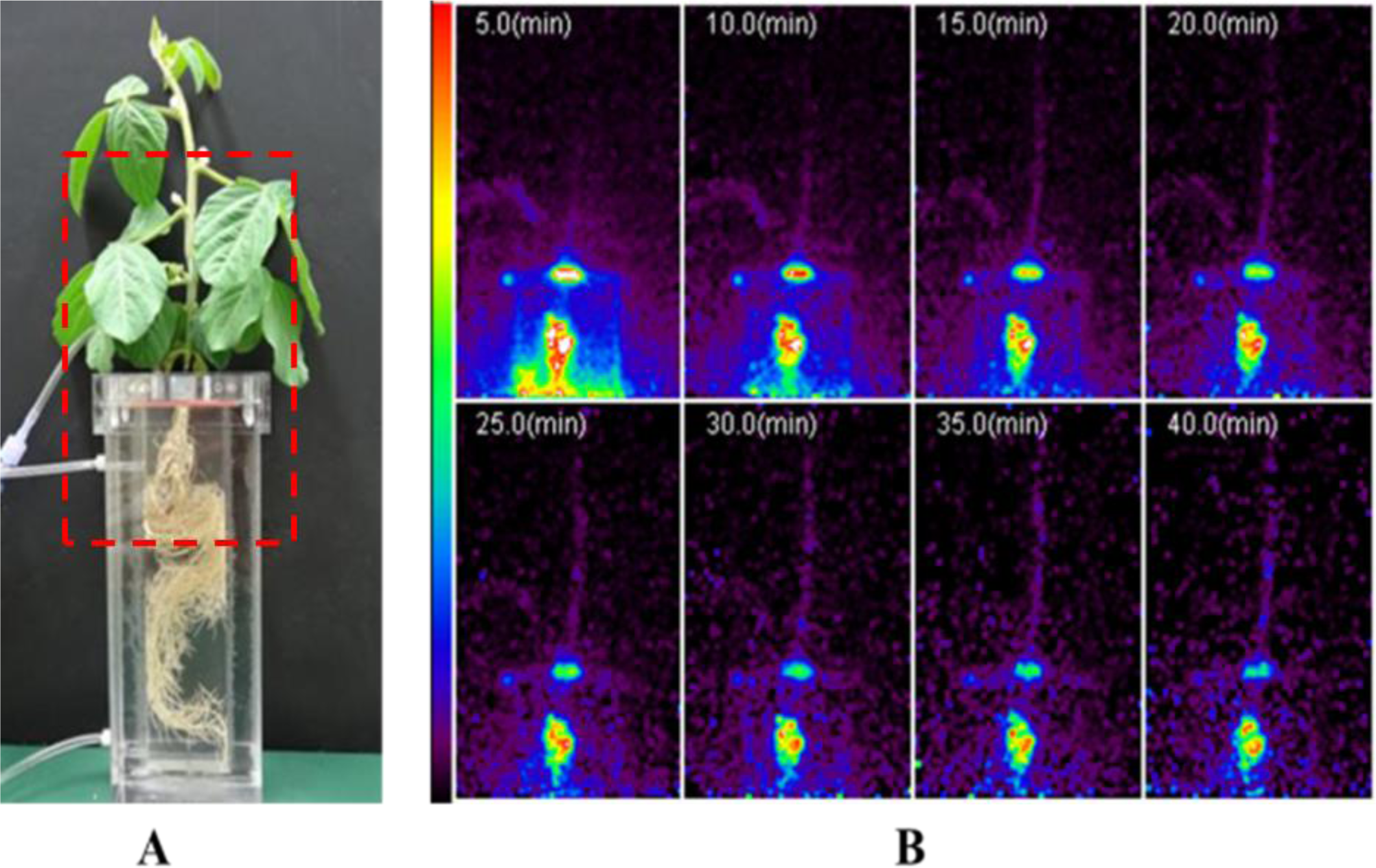
The test plant and Serial PETIS images of the ^13^N movement in the soybean shoot.

The PETIS data was used to analyze the dynamic of nitrogen fixation activity and the translocation of fixed nitrogen from nodules to shoot. To estimate the nitrogen fixation rate and transport velocity of fixed nitrogen, the regions of interest (ROIs) were set on the nodule zone, base stem (ROI1), and upper stem (ROI2) along the stem (**Figure 2A**). **Figure 2B** shows the time-activity curve from the nodule zone. It was estimated from the value just after flushing out of the tracer and the subsequent slope that the average rate of nitrogen fixation was about 0.538 µmol N_2_ h^-1^ and the export rate of fixed nitrogen from nodules to the other parts were evaluated to be 0.017 µmol N_2_ h^-1^ (**Table 1**). The distribution rate of fixed nitrogen form nodules to base stem (ROI1) and upper stem (ROI2) in the initial time was low, it was estimated about 0.0169µmol N_2_ h^-1^ and 0.0101µmol N_2_ h^-1^ at ROI1 (equal to 3.14% at the base stem) and ROI2, respectively (**Table1**).

**Table 1:**
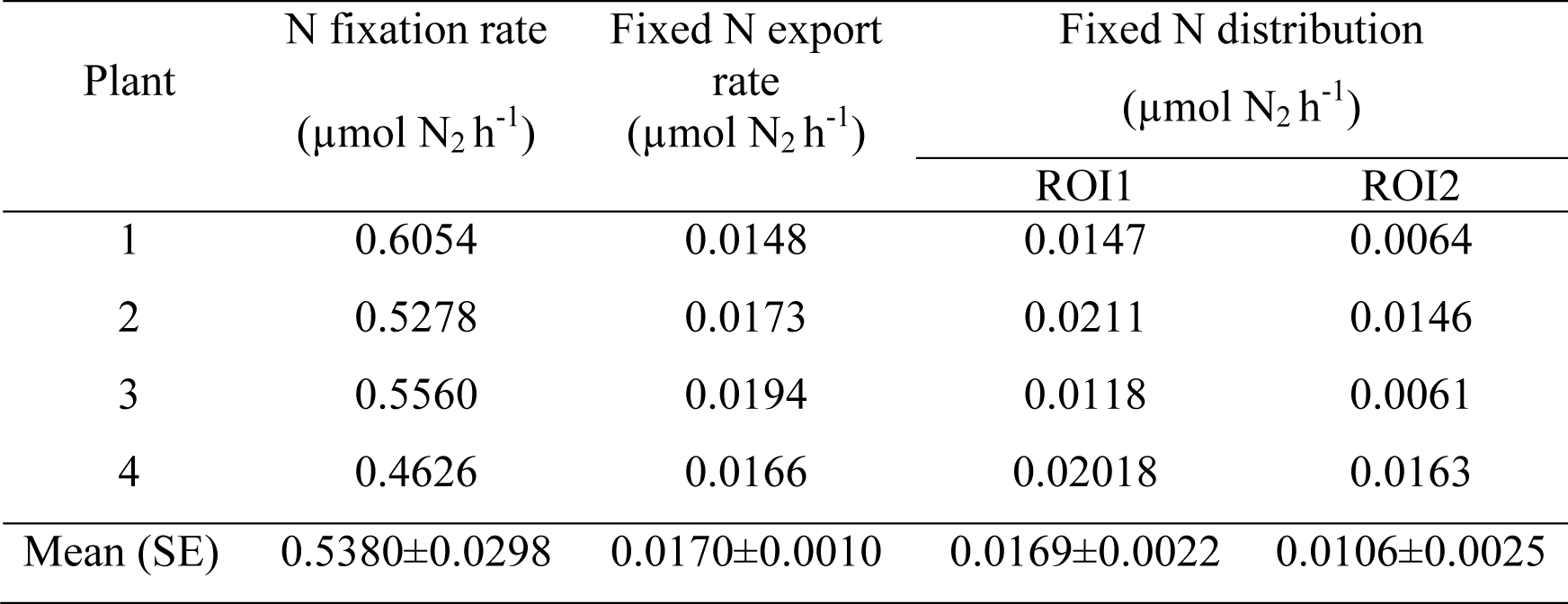
Evaluation of nitrogen fixation rate and export rate of fixed-N (µmol N_2_ h^-1^)

| Plant | N fixation rate<br>($\mu\text{mol N}_2 \text{ h}^{-1}$) | Fixed N export<br>rate<br>($\mu\text{mol N}_2 \text{ h}^{-1}$) | Fixed N distribution<br>($\mu\text{mol N}_2 \text{ h}^{-1}$) | |
| --- | --- | --- | --- | --- |
|  |  |  | ROI1 | ROI2 |
| 1 | 0.6054 | 0.0148 | 0.0147 | 0.0064 |
| 2 | 0.5278 | 0.0173 | 0.0211 | 0.0146 |
| 3 | 0.5560 | 0.0194 | 0.0118 | 0.0061 |
| 4 | 0.4626 | 0.0166 | 0.02018 | 0.0163 |
| Mean (SE) | 0.5380±0.0298 | 0.0170±0.0010 | 0.0169±0.0022 | 0.0106±0.0025 |

To estimate the velocity of the initial transport of fixed-N movement in the stem from root to the shoot, we used the data from TACs at two ROIs to calculate the arrival time of fixed-N at the base stem (ROI1) and the upper stem (ROI2). **Figure 3C** shows the time-activity curves (TACs) from the stem zone (ROI1 and ROI2). From the time-lag between the two TACs, the velocity of movement of fixed nitrogen in the stem was estimated at 1.63 cm min^-1^.

**Figure 3:**
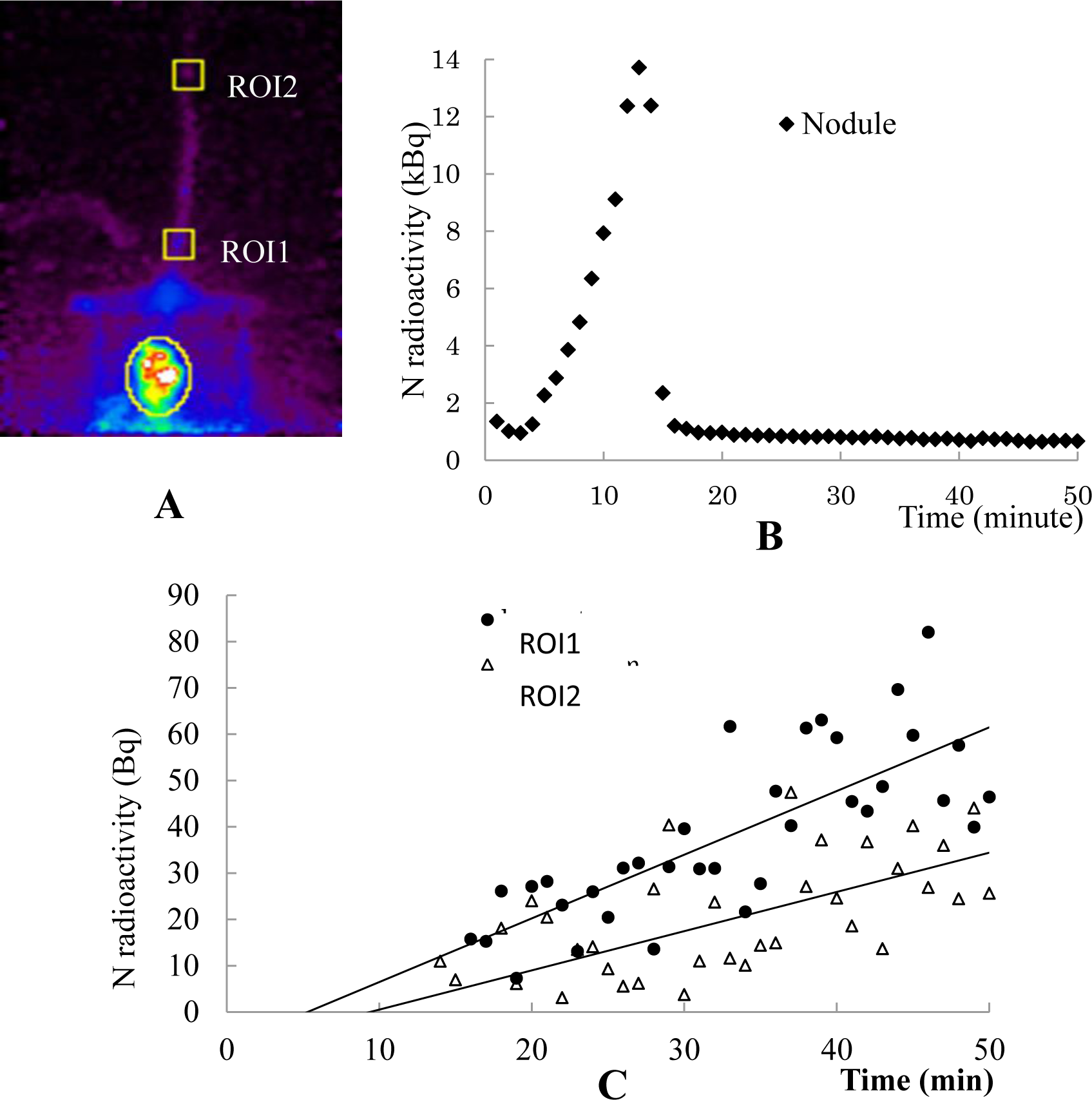
Analysis of time-activity curves generated from PETIS data. A: Two selected ROIs on PETIS image; ROI1 is at the base stem and ROI2 is at the upper stem. B: The time-activity curves showing the fixed N accumulated at nodules. C: The time-activity curves showing the fixed N accumulated at two ROIs

To determine more clearly the transport of fixed nitrogen ([^13^N]N_2_ tracer) in soybean plants we subjected the test plant to the autoradiograph after PETIS investigating. The photograph and BAS image (**Figure 4**) showed the accumulation of fixed-N in nodules and translocation of fixed nitrogen to the shoot. The signal of [^13^N]N_2_ tracer accumulated in nodules was still significantly strong; the radioactivity tracer accumulated in the stem was strong especially at the base stem. This intensity signal was higher in comparison with that of the PEPTIS image. Especially, in BAS image the signal of ^13^N radioactivity was observed in young leaves and new shoot (top bud) but there was no signal in older leaves and roots.

**Figure 4:**
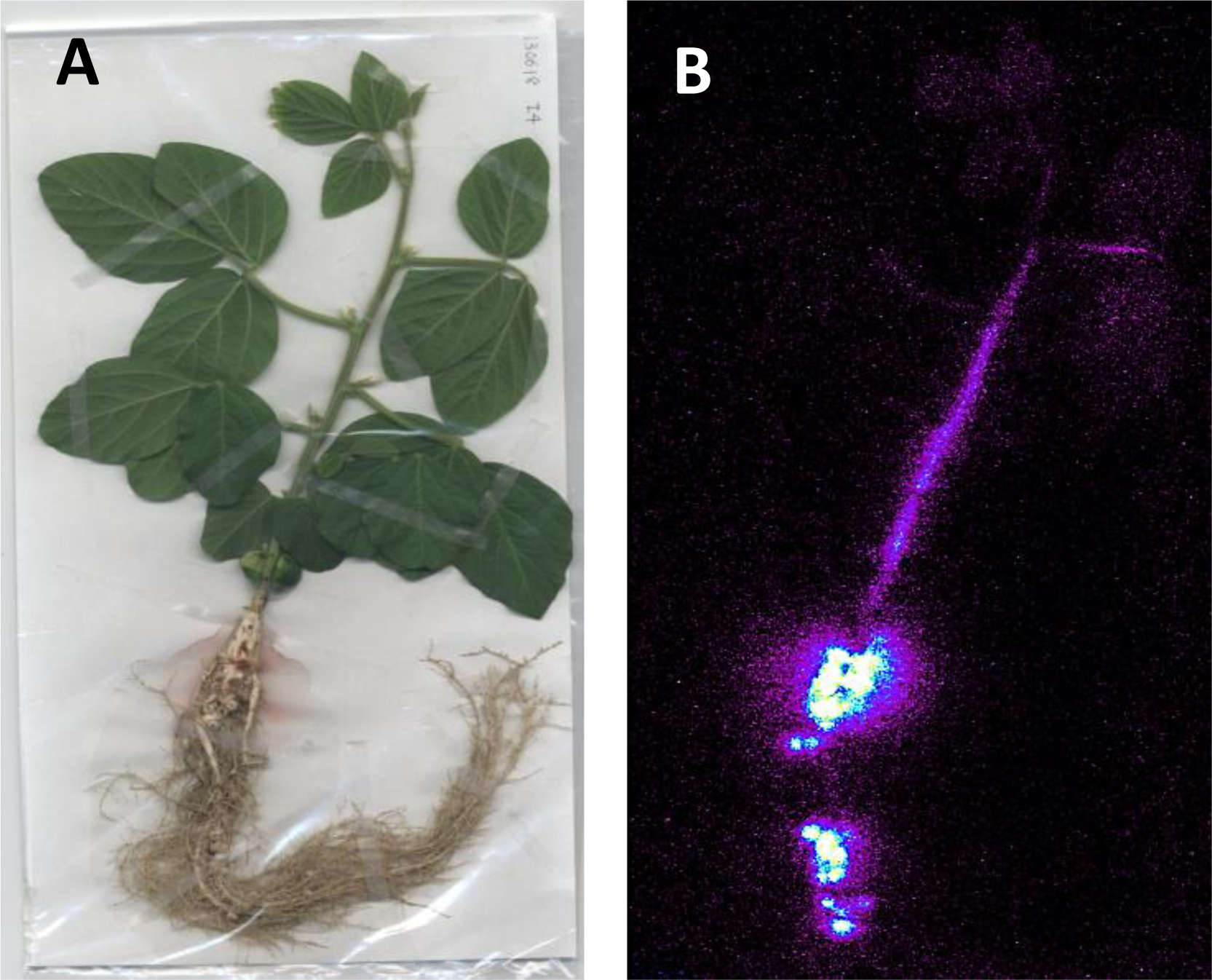
The photograph and autoradiograph of the test soybean plant. **A:** Photograph of soybean plant at 26 DAPs **B:** Autoradiograph

## Discussions

The transportation of fixed nitrogen in legume plants has been studied for a long time, but this mechanism still needs to be elucidated more precisely. By using the PETIS method, the movement of fixed nitrogen in the soybean plant was imaged clearly in this study for the first time. The translocation of fixed nitrogen was observed at base stem at about five minutes after feeding of ^13^N radiotracer and full stem (only in the FOV) at 40 minutes (**Figure 3**). This evidence suggested that fixed-N after synthesized in the nodules, move to the shoot in a short time, which is similar to a previous result that ^13^N was detected first at the trifoliate of soybean plant at 5-10 minutes after introducing ^13^NO_3_^-^ (Sato et al., 1999), and ^13^N reached at base stem of rice in 2 minute and newest leaf in 6 minutes after supplying ^13^NH_4_^+^ (Kiyomiya et al., 2001). It implies that the timing of movement of the fixed-N from soybean nodules was not delayed even if it is compared to that of absorbed nitrogen from the culture solution. In other words, it does not need much time more than a few minutes to convert N_2_ gas into new nitrogen compounds before starting transport of them to others. The velocity of the initial transport of fixed-^13^N was estimated at 1.63 cm min^-1^ at the vegetative stage. This result was similar to previous results found that the speed of ^13^N compound moving in rice plant at vegetative stage was 8.6 cm min^-1^ (Kiyomiya et al., 2001), and the movement of ^13^N fixation compounds, ^13^NO_3_^-^ and ^13^NH_4_^+^ at rate of 6-12 cm min^-1^ in alfalfa root and shoot (Cadwell et al., 1984). Therefore, it was suggested that most of the fixed-^13^N is transported smoothly on the transpiration stream in the xylem from root to shoot as well as nitrate and ammonium.

Although the signal intensity of ^13^N radioactivity was observed early at the base stem, it was still weak at the end of PETIS experiment, and the translocation of nitrogen radioactivity could not be seen in the whole plant because of the limitation of the field of view (FOV) by PETIS experiment. However, this phenomenon was observed more clearly by the BAS image (**Figure 4B**) performed after the end of PETIS measurement. In the BAS image, the signal of N radiotracer was presented only in young leaves and buds (**Figure 4B**). This result suggested that fixed-N was transported in priority to upper organs (young leaves and bud) to create new compounds for plant growth rather than transported to mature leaves. This is consistent with previous results (Hung et al., 2013; Tajima et al., 2004) found that fixed-^13^N was exported to young upper parts of shoot especially stem more than lower parts of the shoot.

When observing the transport of [^13^N]NO_3_ in soybean plant by autoradiography, Sato et al. (1999) also found that the radioactive signal of ^13^N radiotracer was high in young and mature trifoliate leaves compared to primary leaves, while the [^13^N]N radiotracer derived from fixation presented here was only observed in young leaves. This suggested that the fixed nitrogen was translocated to young leaves and buds, while nitrogen that absorbed from fertilizer and soil was transported to all shoots especially mature leaves. It should be also noted in the BAS image that no signal was detected in the nodules and root of the distal region although many nodules attached there. These nodules were immersed in the culture solution so that they could not contact [^13^N]N_2_ tracer gas and could not directly fix it. Therefore, this result suggests that the recycling of fixed nitrogen from the shoots to distant parts of roots and nodules via phloem needs longer than 60 minutes.

The most important finding in the BAS image pointed out that we could not observe the ^13^N radioactive signal in the old leaves although they were close to the source of fixed nitrogen (nodules). This evidence strongly confirmed that fixed-N is transported directly to new vegetative organs and this makes us change the concept that the fixed-N translocation through the shoot may not move only in xylem system as the previous concept that the fixed N in nodule is transported through xylem by transpiration stream by mature leaves (Ohyama et al., 2008; Pate et al., 1979a; Pate et al., 1979b), but the fixed N may be transferred from xylem to phloem in the stem. In the case of fixed N transport, the initial transport of fixed N was mainly in the stem and translocated to young leaves and buds via the phloem system.

The new finding in the initial transport of fixed nitrogen of soybean will become the basis for the next study of fixed N transport in legume plants. However, this is the first result found by using the ^13^N radioisotope method so that it is necessary to do more studies to determine where and how is the fixed nitrogen transferred from the xylem system to the phloem system and transported in stem?

## Acknowledgment

We are thankful to Radiotracer Imaging Group, Department of Radiation-Applied Biology, Takasaki Advanced Radiation Research Institute, Quantum Beam Science Research Directorate National Institutes for Quantum and Radiological Science and Technology, Japan for their facilities support and for providing valuable guidance’s concerning project implementation.

## Abbreviations

PETIS: positron-emitting tracer imaging system
ROI: region of interest
FOV: field of view

## References

Hung, N. V. P., Watanabe, S., Ishikawa, S., Ohtake, N., Sueyoshi, K., Sato, T., Ishii, S., Fujimaki, S., and Ohyama, T., 2013. Quantitative analysis of the initial transport of fixed nitrogen in nodulated soybean plants using ^15^N as a tracer. Soil Science and Plant Nutrition 59, 888–895.

Ishii, S., Ito, N. S. S., Ishioka, N. S., Kawachi, N., Ohtake, N., Ohyama, T., and Fujimaki, S., 2009. Real-time imaging of nitrogen fixation in an intact soybean plant with nodules using ^13^N-labeled nitrogen gas. Soil Science and Plant Nutrition 55, 660–666.

Keutgen, N., Matsuhashi, S., Mizuniwa, C., Itoa, T., Fujimura, T., Ishioka, N. S., Watanabea, S., Sekine, T., Uchida, H., and Hashimoto, S., 2002. Transfer function analysis of positron-emitting tracer imaging system (PETIS) data. Applied Radiation and Isotopes 57, 225–233.

Kiser, M. R., Reid, C. D., Crowell, A. S., Phillips, R. P., and Howell, C. R., 2008. Exploring the transport of plant metabolites using positron-emitting radiotracers. HFSP Journal 2, 189–204.

Kume, T., Matsuhashi, S., Shimazu, M., Ito, H., Adachi, T. F. K., Uchida, H., Shigeta, N., Matsuoka, H., Osa, A., and Sekine, T., 1997. Uptake and transport tracer (^18^F) of positron-emitting in plants. Appl. Radiat. lsot. Vol. 48∼ No. 8 48, 1035–1043.

Layzell, D. B., and Larue, T. A., 1982. Modeling C and N transport to developing soybean fruits. Plant Physiol 70, 1290–1298.

Matsuhashi, S., Fujimaki, S., Kawachi, N., Sakamoto, K., Ishioka, N. S., and Kume, T., 2005. Quantitative modeling of photoassimilate flow in an intact plant using the positron-emitting tracer imaging system (PETIS). Soil Sci. Plant Nutr. 61, 417–423.

McClure, P. R., and Israel, D. W., 1979. Transport of nitrogen in the xylem of soybean plants. Plant Physiol 64, 411–416.

Oghoghorie, C. G. O., and Page, J. S., 1972. Exploration of the nitrogen transport system of a nodulated legume using ^15^N. Planta 104, 35–49.

Ohyama, T., Kaushal, T., Ohtake, N., Sueyoshi, K., Sato, T., Nagumo, Y., Takahashi, Y., Ito, S., Nishigaki, T., and Ishii, S., 2008. Nitrogen fixation and metabolism in soybean plants. Nova Science Publisher, Inc. New York, 16–109.

Ohyama, T., Ohtake, N., Sueyoshi, K., Tewari, K., Takahashi, Y., Ito, S., Nishiwaki, T., Nagumo, Y., Ishii, S., and Sato, A., 2009. Nitrogen Fixation and Metabolism in Soybean Plants. Nova Science Publishers, Inc. New York, 1–131.

Pate, J. S., Atkins, C. A., Hamel, K., McNeil, D. L., and Layzell, D. B., 1979. Transport of organic solutes in phloem and xylem of a nodulated legume. Plant Physiol 63, 1082–1088.

Pate, J. S., Atkins, C. A., Rainbird, R. M., and Woo, K. C., 1980. Nitrogen nutrition and xylem transport of nitrogen in ureide-producing grain legumes. Plant Physiol 65, 961–965.

Pate, J. S., McNeil, D. L., and Layzell, D. B., 1979. Modeling the transport and utilization of carbon and nitrogen in a nodulated legume. Plant Physiol 63, 730–737.

Tajima, S., Nomura, M., and Kouchi, H., 2004. Ureide biosynthesis in legume nodules. Frontiers in Bioscience 9, 1374–1381.

